# MicroRNA-196a is regulated by ER and is a prognostic biomarker in ER+ Breast Cancer

**DOI:** 10.1101/329227

**Authors:** Michael J.G. Milevskiy, Udai Gujral, Carolina Del Lama Marques, Andrew Stone, Korinne Northwood, Lez J. Burke, Julia M.W. Gee, Kenneth Nephew, Susan Clark, Melissa A. Brown

## Abstract

MicroRNAs are potent post-transcriptional regulators involved in all hallmarks of cancer. *Mir-196a* is transcribed from two loci and has been implicated in a wide range of developmental and pathogenic processes, with targets including Hox, Fox, Cdk inhibitors and annexins. Genetic variants and altered expression of *miR196a* are associated with risk and progression of multiple cancers including breast cancer, however little is known about the regulation of the genes encoding this miRNA, nor the impact of variants therein. Here we demonstrate that *MIR196A* displays complex and dynamic expression patterns, in part controlled by long range transcriptional regulation between promoter and enhancer elements bound by ERα. Expression of this miRNA is significantly increased in models of hormone receptor positive disease resistance. The expression of *MIR196A* also proves to be a robust prognostic factor for patients with advanced and post-menopausal ER+ disease. This work sheds light on the normal and abnormal regulation of *MIR196A* and provides a novel stratification method for therapeutically resistant breast cancer.

## Introduction

MicroRNAs are short non-coding RNAs that post-transcriptionally regulate gene expression (1). MicroRNAs have been implicated in many disease, ranging from rare inherited syndromes arising from germline mutations in MiRNA genes through cancers arising from an accumulation of germline and somatic mutations and epigenetic deregulation (2). Research into the biology and pathology of these molecules has led to the identification of clinically useful genetic and epigenetic biomarkers and novel therapeutic agents, often based on antagomiR technology, that have shown promise in the control of disease symptoms and progression (3).

*MicroRNA-196A (miR-196a, MIR196A)* is transcribed in two genomic locations, the *HOXC* (Chr12 in humans, gene *MIR196A2*) and *HOXB* (Chr17 in humans, gene *MIR196A1*) loci, downstream of *HOXC10* and upstream of *HOXB9* respectively. It has been strongly implicated in a range of cancers, primarily as an oncogene. For example, *MIR196A* is overexpressed in breast tumours (4), and a single nucleotide polymorphism (SNP, rs116149130) within the *MIR196A2* gene is associated with a decreased risk of breast cancer (5). *MIR196A* has been shown to target the 3’ UTR of Annexin-1 (*ANXA1*), an important mediator of apoptosis in various pathways (6), in response to the pro-angiogenic vascular endothelium growth factor (VEGF), leading to alterations in angiogenesis. A separate study demonstrated that *MIR196A* could increase growth, migration and invasion of a non-small cell lung cancer cell line through direct targeting of *HOXA5* (7). Two studies have recently shown that *MIR196A* can directly influence the cell cycle by targeting p27/Kip1, an inhibitor of cell cycle progression, to dramatically increase growth and pro-oncogenic features of cancer cell lines (8, 9). Despite the clear importance on *miR-196a* in cancer, its transcriptional regulation remains poorly understood.

Transcriptional regulation is a complex multi-faceted biological process that is significantly altered in cancer. MicroRNA genes are regulated transcriptionally in a similar manner to protein coding and long non-coding RNA genes. Promoters mostly lie upstream (within 10kb of the mature miRNA), contain a CpG island and in an active state when the miRNAs are transcribed by RNA Pol II are enriched for H3K4me3 and lack H3K27me3 similar to protein coding genes (10, 11). Taken together, these data indicate that potential promoters for miRNAs can be identified in a similar manner to methods for protein coding genes. Several instances of miRNA regulation by enhancers have been described, but this area is very much in its infancy (12, 13).

In this study, we aimed to characterise the expression landscape of *MIR196A* including factors regulating its expression and explore potential roles of regulatory elements and factors in breast cancer prognostication.

## Results

### MIR196A expression correlates with HOXC genes in breast cancer

Several *HOXC* protein coding and non-coding genes have shown associations with breast cancer progression. We first identified expression patterns of *HOXC* genes in breast cancers (Supp Figure 1). These data indicate that *MIR196A* expression highly correlates to *HOXC* genes, particularly *HOXC10*.

**Figure 1:**
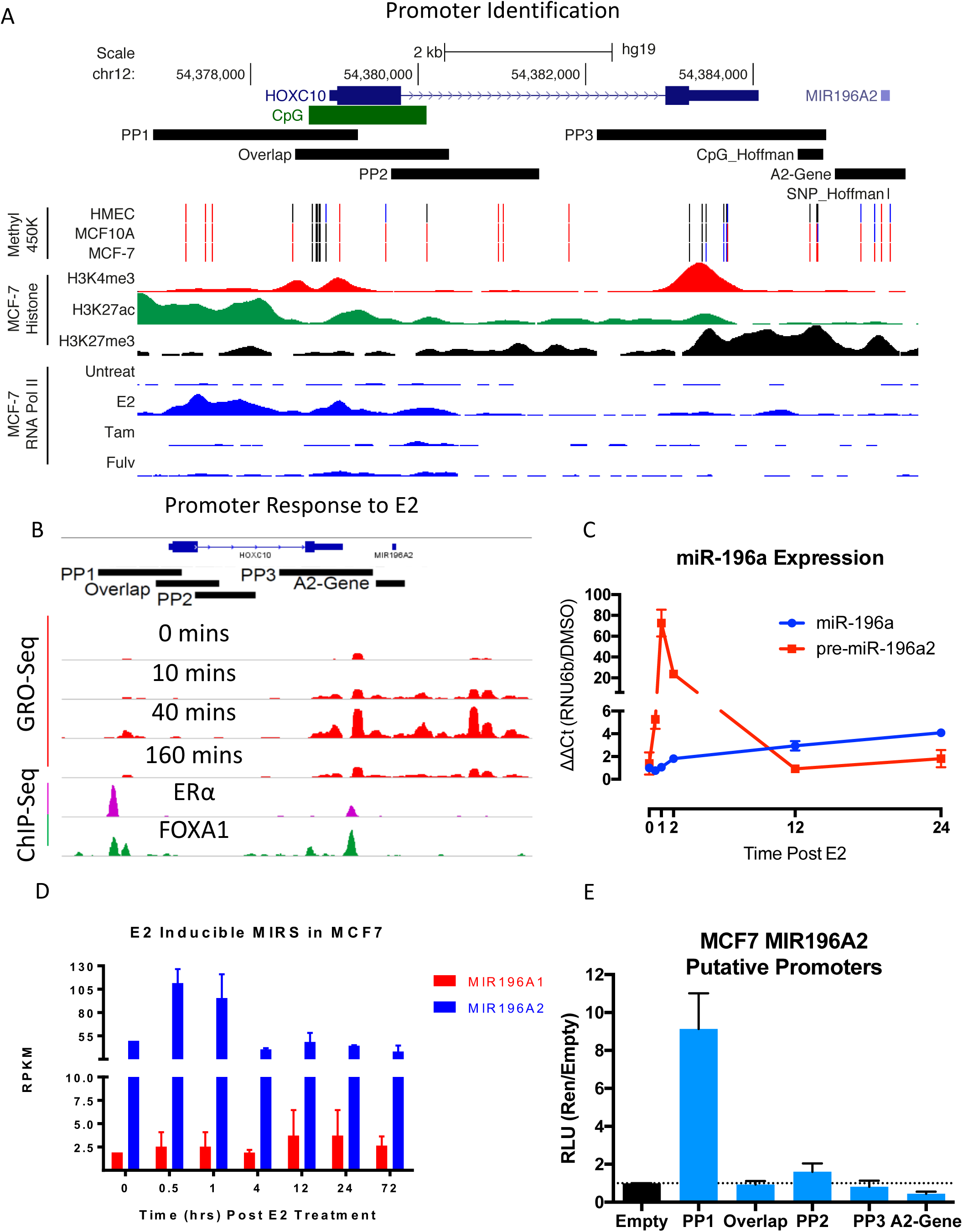
E2 influences *MIR196A2* expression in breast cancer. A) Identification of putative promoter regions for the *MIR196A2* gene using histone markers and ChIP-Seq indicated in figure (E2 = oestradiol, Tam = 4-hydroxytamoxifen, Fulv = Fulvestrant). Refseq genes are indicated in blue at the top with coordinates based on hg19 chromosome 12. Putative promoter regions (PP1,2,3) and the previously implicated SNP (rs116149130) and CpG from Hoffman *et al* (5) are indicated by black rectangles. MCF7 DNA methylation 450K array data indicate unmethylated (black), partial methylation (blue) and methylated (red). **B)** GRO-Seq measurements of RNA Polymerase engagement and elongation points from the putative promoters, after E2 stimulation in MCF7 cells. Lower part, ChIP-Seq for binding of ERα and FOXA1 to the putative promoters. **C)** qRT-PCR the MIR196A2 response to E2 in MCF7 cells. Qiagen precursor primers were used to detect the precursor miRNA at the specified time points and CT values were normalised to a DMSO vehicle control and the qPCR control of *RNU6b*. **D)** MiRNA-Seq reads per kilobase per million (RPKM) for the precursor miRNAs following E2 addition to MCF7 cells. **E)** Luciferase reporter assay measuring the influence of *MIR196A2* putative promoter on the luciferase gene transcription. Measurements are relative light units (RLU) normalised to the renilla plasmid (pRL-TK) acting as a transfection control and to the pGL3/Empty plasmid. Experimental measures are done in triplicate with the experiment repeated, data not shown.

Next we investigated whether these associations are also observed in normal cells of the human breast. The association between *MIR196A* expression and *HOXC* genes is more limited, observed most strongly with *HOXC11* and *HOXC10*, the genes surrounding the *MIR196A* gene (Supp Figure 2A). *MIR196A* appears to be mostly expressed within the basal stem-cell (BSC) derived cells, whilst much lower in expression of the more differentiated cell types (Supp Figure 2B).

**Figure 2:**
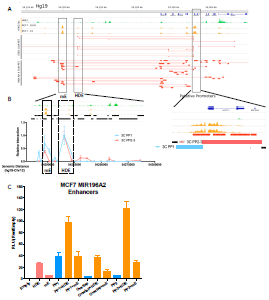
Distal putative enhancer elements of the MIR196A2 putative promoters. A) Histone modification and ChIA-PET for ESR1 and RNA Pol II in MCF7 cells of the HOXC locus and corresponding gene desert. The histone modification H3K27ac is a measure of regulatory element activity and was assessed in HMEC and MCF7 cells plus or minus E2. ChIA-PET interactions are represented by red lines, solid rectangles indicate the sequenced tag and the two points that were physically interacting and tethered to either ESR1 or RNA Pol II. **B)** Zoom in of the *MIR196A2* enhancer (mE) and *HOTAIR* distal enhancer (HDE) (left) and the putative promoter elements (right). Black rectangles indicate the genome fragment sizes post digestion with *Hind*III. Graph is 3C-qPCR for either the 3C PP1 or 3C PP2-3 fragments with the Y-axis the relative interaction and the X-axis the genomic location. All genomic coordinates were based on chromosome 12 in the hg19. C) Luciferase reporter assay showing the augmentation of HOXC promoters with either mE or HDE, shown a RLU normalised to the co-transfected control renilla plasmid and to the vector backbone pGL3-Basic.

### MIR196A expression is regulated by oestrogen

We and others have previously demonstrated regulation of *HOXC* genes by oestrogen in breast cancer (14-18). Given that *MIR196A* expression strongly correlates with expression of HOXC protein coding genes in breast cancer (Supp Figure 1), we sought to determine if oestrogen also regulates *MIR196A2.* Chromatin immunoprecipitation (ChIP-Seq) for RNA polymerase II demonstrates that polymerase binding in the region surrounding the *HOXC10* gene and *MIR196A* gene is dependent on oestrogen in MCF7 cells and is repressed with both tamoxifen or fulvestrant treatment (Figure 1A). Global-run-on sequencing (GRO-Seq) is able to measure nascent RNA, assessing changes in transcription with high sensitivity. Analysis of MCF7 GRO-Seq data clearly indicates a dramatic increase in RNA production in the genomic region surrounding *MIR196A2*, peaking at 40 mins following addition of oestradiol (E2) (Figure 1B). This increase in RNA production from the *HOXC* locus was validated with qRT-PCR and RNA-Seq from MCF7 cells following addition of E2 (Figures 1C and D). These data clearly indicate an increase in precursor miRNA from *MIR196A2* but not *MIR196A1* in response to E2. Taken together this suggests that *MIR196A2* is transcriptionally regulated by oestrogen.

### Transcriptional regulation of miR196A

To identify the structural elements associated with the transcriptional regulation of *MIR196A*, histone methylation patterns in the MCF7 breast cancer cell line were assessed. This analysis uncovered putative promoter elements upstream of *MIR196A* including a shared promoter with *HOXC10* (Figure 1A).

Given that *MIR196A* expression is regulated by oestrogen we hypothesized that its transcription may be controlled by the oestrogen receptor (ER). Using publically available datasets we established that oestrogen mediated upregulation of *miR-196A* expression is accompanied by binding of ERα and its pioneer factor FOXA1 to two putative promoter regions, putative promoters 1 and 3 (PP1 and PP3), upstream of the miR196A2 transcription start site (Figure 1B).

Upon cloning of these putative promoter sites into luciferase reporter vectors where PP1 and also PP2; modestly; increases luciferase gene transcription (Figure 1E), with the most active promoter in MCF7 cells, PP1 (*HOXC10* promoter).

Given that ERα often binds to distal enhancer elements to exert its function, we examined the hypothesis that *MIR196A2* is controlled by long-range transcriptional regulation, mediated by ERα tethered gene looping. Using ChIA-PET (Chromatin Interact Analysis by Paired End Tags) genome-wide chromatin interactions that immunoprecipitate with either ERα or RNA Polymerase II (correlative with active promoters and enhancers), we identified two major sites of interaction with the *MIR196A2* promoters (Figure 2A). One of these is a previously identified HOTAIR enhancer (HOTAIR distal enhancer, HDE (15)) and the other a novel interacting partner (MIR196A2-Enhancer, mE). Chromosome conformation capture (3C) digestion of the HOXC genomic locus digests the *MIR196A2* region into two fragments. 3C-qPCR demonstrates that both enhancer elements physically interact with each of the two *MIR196A2*/*HOXC10* promoter region (Figure 2B). Cloning of these fragments downstream of the putative promoter luciferase reporters clearly demonstrates significant augmentation of transcription for both the PP1 and PP2, with HDE appearing to be the most active in MCF7 cells (Figure 2C).

Interestingly, a previous study (5) identified a SNP and an upstream CpG island associated with a decrease in breast cancer risk. This SNP lies within the *MIR196A2* gene and the CpG island (CpG_Hoffman) is immediately upstream, falling into the 3’ end of the PP3. Analysis of DNA methylation reveals that this CpG island is mostly methylated in non-malignant MCF10A and cancerous MCF7 cells, whilst unmethylated in human mammary epithelial cells (HMEC) (Figure 1A).

### MIR196A is differentially expressed in breast cancer

Given that *MIR196A* is regulated by ERα, we investigated its expression patterns in relation to commonly utilised molecular markers of breast tumours (Figure 3A). This analysis identified four distinct clusters of *MIR196A* expression (Clusters 1-4). Interestingly clusters 1 and 3 show a strong correlation to expression of hormone receptors (HR) (*AR, ERα, PGR, HER2*) and HR cofactors (Figure 3B). In contrast, clusters 2 and 4 have significant negative correlation to expression of *ERα, PGR, FOXA1* and *GATA3*, whilst associating with *EGFR* and *HER2*. This expression is further defined by the PAM50 intrinsic subtypes where *MIR196A* is strongly expressed in the HER2 subtype, whist in the luminal A and B subtypes expression is very dynamic (Figure 3C).

**Figure 3:**
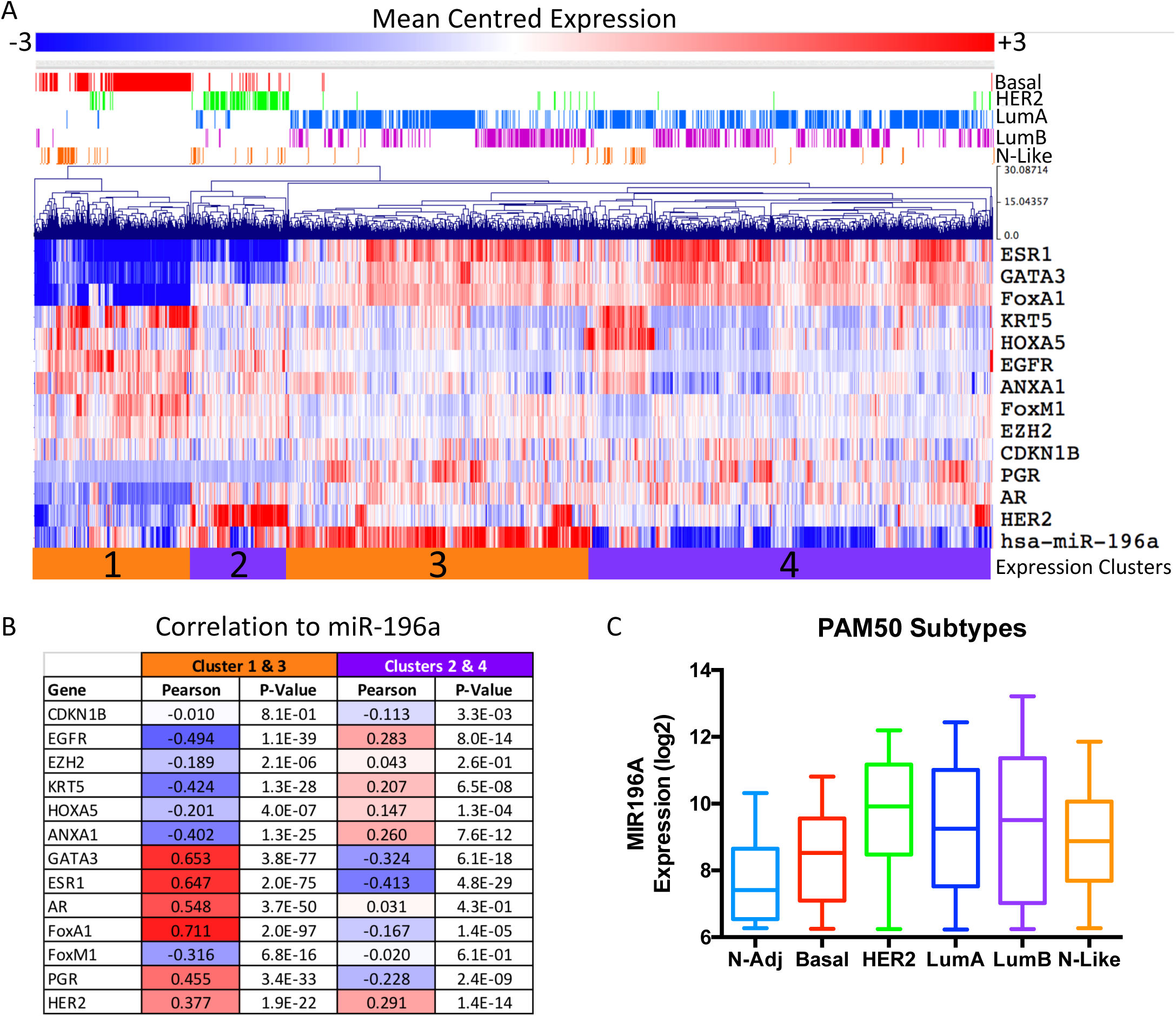
*MIR196A* is differentially expressed in breast cancer. A) Mean-centred log2-expression of *MIR196A* and commonly utilised breast cancer molecular markers. Expression values were hierarchically clustered and the PAM50 tumour subtypes are indicated above the plot. Expression values are indicated by colour scale bar. **B)** Pearson correlation coefficients, and corresponding P-values, for each gene against the expression of *MIR196A* either in the orange or purple clusters. **C)** Intensity values for the expression of *MIR196A* across the five molecular subtypes, PAM50. All data was sourced by the METABRIC cohort (61, 62).

### MIR196A is a biomarker of breast cancer progression

To further explore the expression of *MIR196A* in breast cancer, we utilised expression data from the METABRIC cohort of breast tumours. Expression analysis of this miRNA indicate that it is significantly over-expressed in breast tumours compared to normal adjacent tissue and over-expression is associated with an increase in tumour stage (Figure 4A and 4B). Interestingly, high expression of *MIR196A* is associated with a poor survival in estrogen receptor positive (ER+) breast cancer, whilst high expression associates with a better outcome in triple-negative breast cancer (TNBC) (Figure 4C and D).

**Figure 4:**
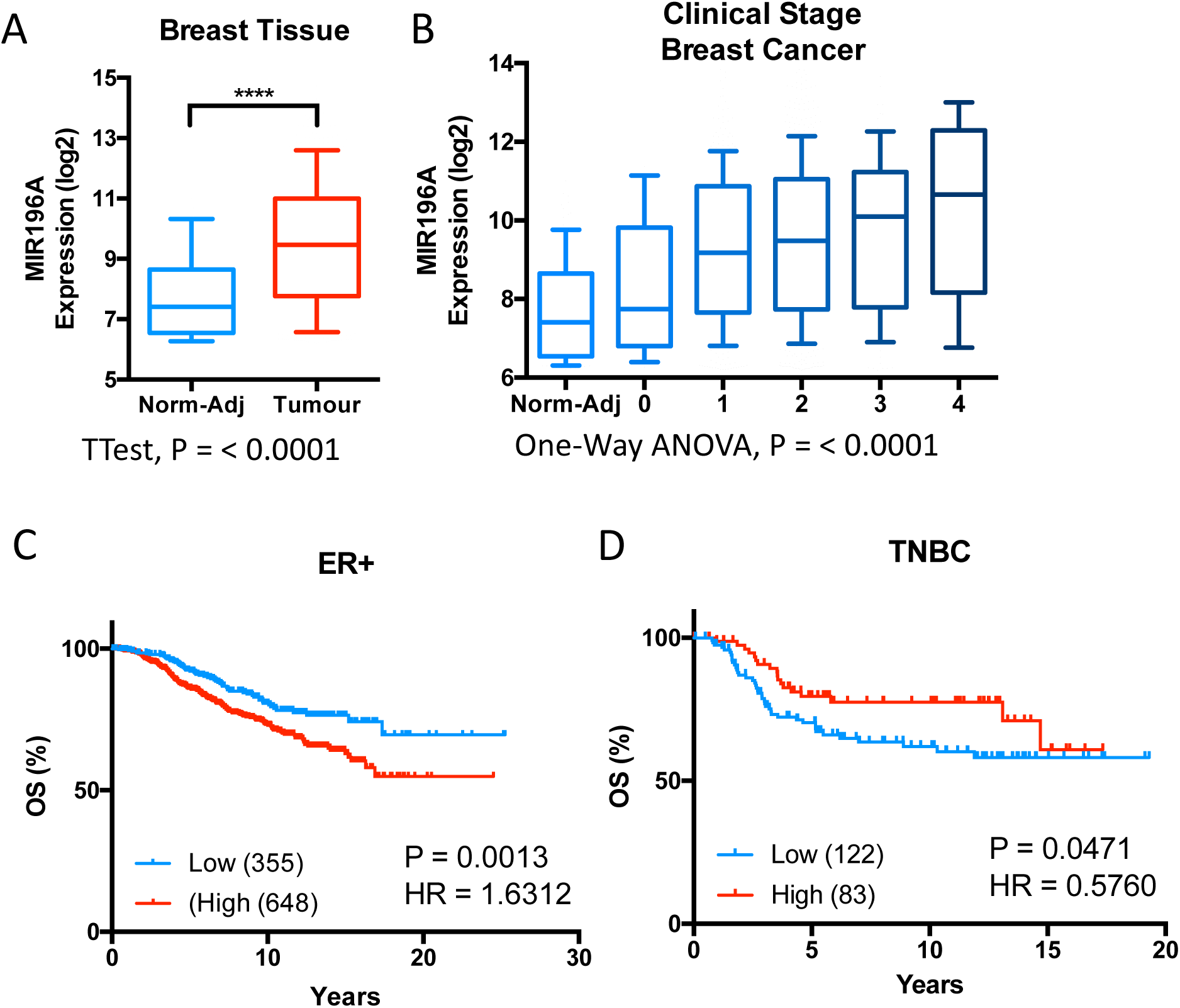
*MIR196A* is a biomarker of breast cancer progression. A) Log2 miR-Array intensity for the expression of *MIR196A* in normal adjacent tissue and breast tumours. A T test was used to find statistical significance with **** = a p value of < 0.0001. **B)** Log2 intensity for the expression of *MIR196A* in normal adjacent and tumour stages 0 to 4. A One-Way ANOVA was used to find a significant trend with a p-value = < 0.0001. **C and D)** Kaplan-Meier curves stratifying overall survival of breast tumours by expression of *MIR196A*. Log-rank p-value (P) and hazard ratios (HR) displayed. All expression data sourced from METABRIC (61, 62).

### MIR196A is a prognostic biomarker in advanced ER+ disease

Using *MIR196A* expression, overall survival of ER+ tumours responding to both hormone therapy (HT) and chemotherapy (CT) was stratified (Figure 5A). Women with low *MIR196A* expression had exhibited a high rate of survival (>95% at 10 years), whilst most women within the high expression group died within 17 years (61% at 10 years).

**Figure 5:**
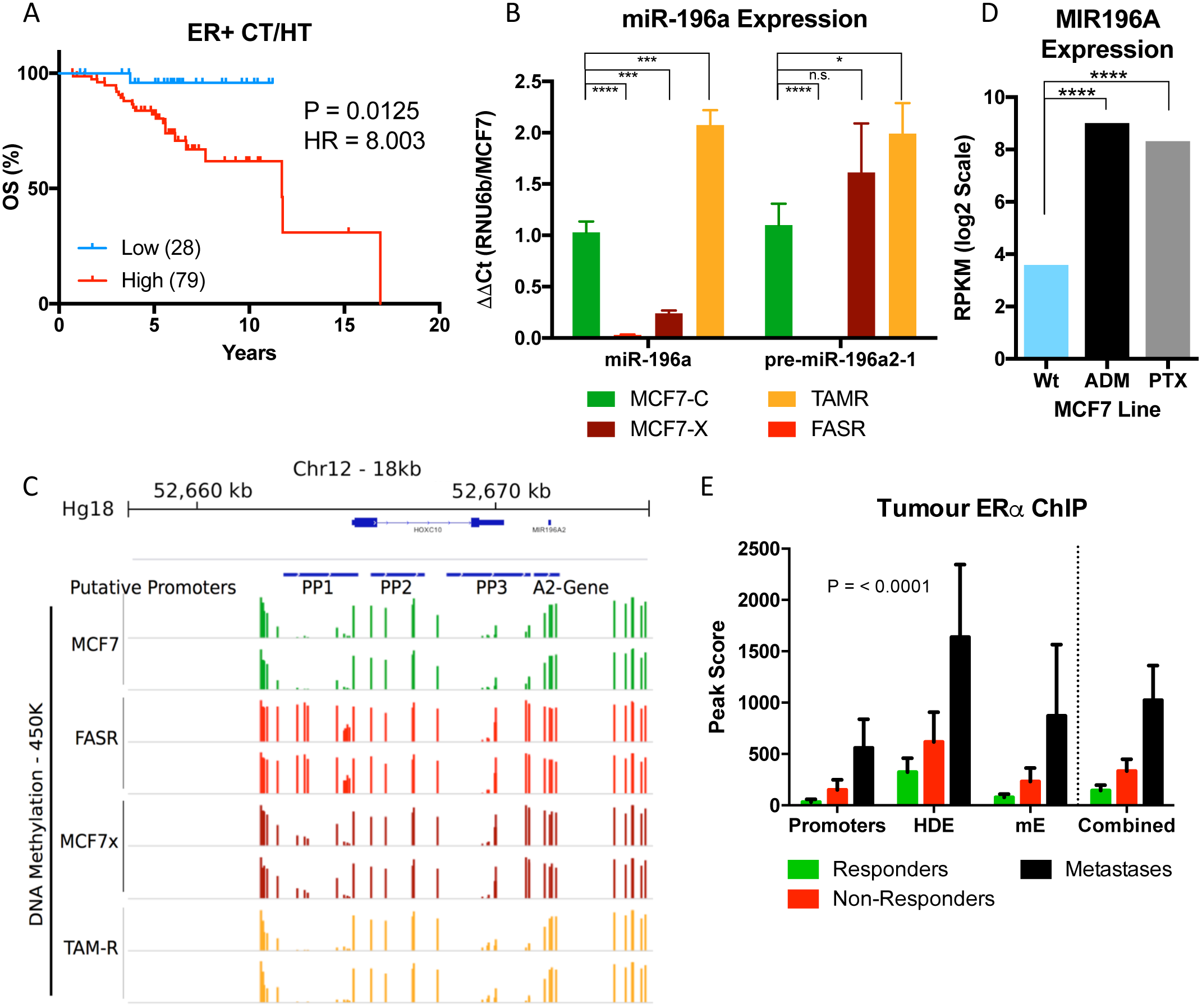
Therapeutic resistance leads to increase in *MIR196A* expression. A) qRT-PCR relative expression for the mature and precursor *MIR196A* transcripts in MCF7 derived cell line models of endocrine therapy resistance. miRNA expression values are normalised to the expression of *RNU6b* and the MCF7 cell line. Error bars are the standard deviation of two technical replicates and four biological replicates. MCF7-C = Control, MCF7-X = Oestrogen deprived, TAMR =Tamoxifen resistant and FASR =Fulvestrant resistant. **B)** Corresponding DNA methylation for MCF7 derived cell lines, as measured by 450K methylation array, for the *MIR196A2* genomic region. **C)** Log2 reads-per-kilobase-per-million (RPKM) expression of *MIR196A* in MCF7 wild-type and adriamycin (ADM, aka Doxorubicin) and paclitaxel (PTX) derived resistance cell lines. **D)** Peak scores for the binding of ERα to *MIR196A2* regulatory elements in ER+ breast tumours. Peak scores were generated using MACS, normalised to the Input control for the ChIP-Seq library. Peak scores are the average for 9 responders, 9 non-responders and 3 metastases. Data is sourced from Ross-Innes *et al* (55). **E)** Kaplan-Meier survival curves for patients with ER+ disease, treated with both chemotherapy (CT) and hormone therapy (HT). Patient survival is stratified by expression of *MIR196A* into low or high expression subgroups. Expression and survival data sourced from METABRIC (61, 62).

Given that *MIR196A* is regulated in part by oestrogen, and the disparity in prognostication of ER+ and TNBC, we investigated the effects of menopause on the stratification of survival for ER+ women. The effects of menopause on the human breast are largely unknown, however serum levels of oestrogen and progesterone dramatically reduce post menopause. In pre-menopausal women, high expression of *MIR196A* is associated with a good outcome in ER+ disease (Table 1). Multivariate analysis demonstrates that *MIR196A* is one of the few significant biomarkers for ER+ tumours arising before menopause. In post-menopausal women, all tested biomarkers were significant in ER+ disease, including *MIR196A*, however high expression is now associated with a poor outcome.

**Table 1:** Menopause effects the stratification of patient survival by *MIR196A* in ER+ disease. The overall survival of patients with ER+ disease was stratified by *MIR196A*, HER2, PGR expression or commonly utilised clinical markers. On the left is the univariatie cox-proportional hazard ratios for each condition, and the right the multivariate cox-proportional hazard model and the conditions which contribute to the most significant model. HR = Hazard ratio, CI = Confidence interval. Expression and survival data sourced from METABRIC (61, 62).

| ER+ Pre-Menopausal |  |  |  |  |  |  |
| --- | --- | --- | --- | --- | --- | --- |
| Condition | Univariate Cox-proportional hazard |  |  | Multivariate Cox-proportional |  |  |
|  | HR | (95% CI) | P-Value | HR | (95% CI) | P-Value |
| HER2 (high vs low) | 2.695 | 1.2479 - 5.8213 | 0.0120 | 3.352 | 1.3483 - 8.3325 | 0.0096 |
| MIR196A (high vs low) | 0.463 | 0.2325 - 0.9202 | 0.0288 | 0.342 | 0.1534 - 0.7623 | 0.0091 |
| Tumour Grade (1,2,3) | 1.638 | 0.9988 - 2.6844 | 0.0517 |  |  |  |
| Tumour Stage (0-4) | 1.652 | 0.9937 - 2.7472 | 0.0541 |  |  |  |
| Lymph Node (+, -) | 1.857 | 0.9836 - 3.5069 | 0.0575 |  |  |  |
| Size (T1, T2, T3) | 1.470 | 0.9518 - 2.2688 | 0.0840 | 1.798 | 1.0893 - 2.9693 | 0.0225 |
| PGR (high vs low) | 0.552 | 0.2770 - 1.0978 | 0.0919 |  |  |  |
| Age At Diagnosis | 0.985 | 0.9261 - 1.0469 | 0.6221 |  |  |  |
| ER+ Post-Menopausal |  |  |  |  |  |  |
| Lymph Node (+, -) | 2.739 | 2.0075 - 3.7358 | <0.0001 | 1.720 | 1.1510 - 2.5711 | 0.0085 |
| Tumour Stage (0-4) | 2.363 | 1.8658 - 2.9926 | <0.0001 | 1.519 | 1.0557 - 2.1868 | 0.0251 |
| Size (T1, T2, T3) | 1.866 | 1.4837 - 2.3461 | <0.0001 | 1.560 | 1.1146 - 2.1832 | 0.0099 |
| Tumour Grade (1,2,3) | 1.822 | 1.4057 - 2.3622 | <0.0001 | 1.455 | 1.0926 - 1.9385 | 0.0107 |
| MIR196A (high vs low) | 1.847 | 1.3065 - 2.6110 | 0.0005 | 1.599 | 1.0806 - 2.3652 | 0.0195 |
| HER2 (high vs low) | 2.165 | 1.3982 - 3.3521 | 0.0006 | 2.210 | 1.3624 - 3.5847 | 0.0014 |
| PGR (high vs low) | 0.636 | 0.4708 - 0.8594 | 0.0034 |  |  |  |
| Age At Diagnosis | 1.023 | 1.0059 - 1.0403 | 0.0086 |  |  |  |

### Therapeutic resistance leads to increases in MIR196A expression

TNBC is resistant to hormone-based therapies and HR+ disease often becomes resistant to anti-oestrogen treatment. Using established models of HR+ disease resistance we found that *MIR196A* expression is significantly increased in tamoxifen resistant MCF7 cells (TAMR) whilst it is almost depleted in fulvestrant resistance (FASR) (Figure 5B). These expression patterns match changes in DNA methylation to the *HOXC10*/*MIR196A2* promoters in these same cells (Figure 5C). For HR+ resistant tumours the only remaining therapeutic options are radiotherapy and chemotherapy. Using RNA-Seq data for cell line models of resistance to paclitaxel and adromycin, two common chemotherapeutics, *MIR196A* expression again increases in resistant cell lines compared to the treatment sensitive cell line (Figure 5D).

Utilising ERα ChIP-Seq performed in human patients with HR+ disease, binding sites for ERα were identified in the genomic region of *MIR196A*. This tumour cohort contains women who respond to HR therapy, those who do not and metastases from resistant tumours. An increase in ERα occupancy is seen at both enhancer and promoter regions of *MIR196A* in non-responders and metastases (Figure 5E). The increased genome-wide ERα binding in the more resistant tumours was shown by the authors to associate with changes to expression patterns crucial for the resistant tumour to survive therapy and become resistant.

## Discussion

The expression of *MIR196A* in breast cancer is both dynamic and complex. In this paper, we have elucidated important elements, factors and mechanisms controlling the transcriptional regulation of *MIR196A* and shown that changes in this regulation are associated with breast cancer progression and therapeutic resistance.

Previous genetic association studies have shown that SNPs within *MIR196A2* confer a reduced risk of breast cancer (5, 19, 20). Hoffman and colleagues (5) postulated that the polymorphism located within the *MIR196A2* gene reduces microRNA maturation thereby reducing expression of the gene. They also identified that an upstream CpG island is associated with reduced risk when hypermethylated. Here we show that this upstream CpG island lies within the transcriptionally active region of *HOXC10* and *MIR196A2* as observed through GRO-Seq. Interestingly, this CpG island is completely methylated in models of oestrogen deprivation and fulvestrant treatment, but not in tamoxifen resistant cells. DNA methylation is most commonly associated with repressed transcription (21), hypermethylation of this region in a transcriptional high region may severely impair expression. Given that various transcription factors strongly influence transcription in endocrine resistant breast cancer, these data suggest that binding of ERα accompanied by cofactors may be needed to maintain low methylation levels and active transcription in breast cancer (22-26).

We have previously demonstrated that long-range regulation of *HOXC* genes occurs in breast cancer and is influenced by ERα and its associated cofactors (15). *HOX* gene expression is tightly controlled in a spatiotemporal manner to ensure proper axial formation along the anterior-posterior axis (27). Within the cell types of the human breast, *HOX* gene expression appears dynamic and the association between *MIR196A* and *HOXC* genes is not significant. The strong correlation in expression of all *HOXC* genes in breast tumours with *MIR196A* is in stark contrast to expression in normal tissues. Several instances have been described regarding the influence of multiple distal enhancers on gene expression, such as the well characterised locus-control-region (LCR) of the Beta-globin genes or the c-Myc enhancers active across multiple cancer types (28-31). Given the extensive interactions between this locus and its adjacent gene desert, we hypothesise that a consorted effort of multiple enhancers is responsible for the overexpression of these genes in cancer possibly driven by extensive binding and activity of ERα. To explore this hypothesis a high resolution chromatin interaction analysis of this region in breast cancer cells would be required, such as 5C (32) or NG Capture-C (33), coupled with ERα ChIP-Seq and ChIA-PET (34).

Whilst this manuscript was in preparation new data has come to light which corroborates our conclusions. Jiang *et al* (35) demonstrate that the mature *MIR196A* transcript positively responds to oestrogen stimulation in MCF7 cells, and this is mediated by upstream ERα binding. This binding peak falls within PP3. Whilst we show that PP3 is not able to increase luciferase expression in a luciferase reporter assay, the binding of ERα may be important for the activity of the *HOXC10* and *MIR196A2* promoters.

Using hierarchical clustering of breast tumour RNA-Seq data, we observed two distinct expression patterns associated with *MIR196A* expression. Data presented here suggests that the two loci encoding for this miRNA contribute greatly to the complexity of its expression. It is currently unclear how dual-encoded miRs are regulated of which many exist (36).

Interestingly, DNA methylation at several sites within the *HOXC* locus negatively correlates with the expression of this miRNA, supporting the notion of DNA methylation as a repressive epigenetic modification in this context (21).

High expression of *MIR196A* is a biomarker of poor prognosis in ER+ tumours, especially in those patients resistant to therapy. Expression of *MIR196A* increases in response to tamoxifen and chemotherapeutic agents in oestrogen responsive MCF7 cells. This increase in expression is associated with loss of DNA methylation within the promoter regions of the miRNA. In poor responders with ER+ tumours, *HOXC* enhancer elements appear to more readily bind the ER. These data raise the possibility that the pathway to resistance to therapy in ER+ tumours involves the de-repression and over-activation of promoter and enhancer elements. This is commonly seen throughout cancer (37-39), with suggestions that enhancer disruption can revert cells to a non-terminally-differentiated state a common hallmark of tumourigenesis. *HOX* genes are essential in embryonic development, these genes would be a valuable asset for any tumour cell to use to sustain a stem-cell like state (40, 41).

Breast cancer incidence and relative subtype changes after menopause (42, 43). In women younger than 45, luminal breast tumours account for 33-44% (44, 45). This increases to 70-72% in women older than 65. In contrast, basal-like tumours are more common in younger women, suggesting a switch or evolution in the factors driving cancer following menopause, most likely related to the decline in oestrogen production. It is then interesting to note that higher expression of *MIR196A* associates with good outcome in pre-menopausal women with ER+ tumours, and a poor outcome of ER+ tumours following menopause. Given the strong involvement of *HOX* genes in development, we hypothesise that there is a change in the regulation and expression of these genes through and following menopause, which in turn impacts their contribution to the development of certain breast cancer subtypes.

*MIR196A* is a dynamically expressed miRNA in both normal mammary cells and breast tumours. This miRNA is a possible biomarker for the progression of breast tumour to becoming resistant to therapy. Future studies should aim to uncover the purpose of increase *MIR196A* expression and if it is required for development of resistance alone or in combination with other *HOXC* genes.

## Material and Methods

### Cell Culture

MCF7 cells, for the development of endocrine resistance sub-lines were obtained from AstraZeneca. MCF7, Tamoxifen-resistant (TAMR), Fulvestrant-resistant (FASR), and oestrogen-deprived (MCF7x) cells were cultured as described (46-48). All cell lines were cultured for less than 6 months after authentication by short-tandem repeat (STR) profiling (Cell Bank, Australia).

### Cloning and reporter assays

All PCR products for luciferase reporter assays were ligated into Invitrogen’s pCR-Blunt plasmid using T4 DNA Ligase, at 4^±^C overnight. *MIR196A* enhancers and promoters were digested from pCR-Blunt and cloned into the luciferase reporter plasmid pGL3-Basic. Enhancers were cloned into the *Bam*HI/*Sal*I site whilst promoters were cloned into the multiple cloning site immediately upstream of the luciferase gene. See Supplementary Table1 for primers.

MCF7 cells were transfected in antibiotic free media with 500 ng of modified pGL3 reporter constructs, 20 ng of pRL-TK (Renilla transfection control) and with 0.5μL of Lipofectamine 3000 (Life Technologies, L3000-008). 48 hours post transfection luciferase readings were measured using a DTX-880 luminometer and Dual-Glo Stop and Glo luciferase reporter kit (Promega, E2920), following the manufacturer’s recommended protocol.

### RNA extraction and Gene Expression

Cell lysates were prepared using Life Technologies TRIzol^®^ reagent and RNA was chloroform extracted and isopropanol precipitated. RNA was DNaseI treated with the DNA free kit from Ambion (Life Technologies, AM1906). RNA for miRNA analysis was reverse transcribed using the miScript RT II kit from Qiagen (218161), following instructions as per the manufacturer. Assays for all miRNAs were performed with Qiagen’s miScript SYBR Green PCR Kit (218073). Primers specific to each mature or precursor miRNA were assayed coupled with a universal primer, see Supplementary Table 2 for assay IDs. Expression data for miRNAs was normalised to the snoRNA RNU6b. All qRT-PCRs were performed using the protocols advised by the manufacturers on a Corbet Rotorgene-6000.

RNA-Seq on MCF7 cells following oestradiol treatment was performed as described previously by K. Nephew (see author list) (49). RNA-Seq from Adriamycin (ADM) and paclitaxel (PTX) resistant MCF7 derived cells was sourced from GSE68815 (50). Expression of HOX genes in human breast cells was sourced from Gascard *et al* (51).

### Genomic Data Analysis

Accession codes for publically available data were, MCF7 ChIP-Seq (GSE14664, (52)), GRO-Seq (GSE27463, (53)), ChIA-PET (GSE39495, (34, 54)), Breast tumour ERα ChIP-Seq (GSE32222, (55)). MCF7 histone ChIP-Seq and breast cell 450K array data was sourced from ENCODE via http://genome.ucsc.edu/ENCODE/downloads.html. ChIP-Seq data was mapped with Bowtie (56) and peaks called by MACS (57) and viewed in the Interactive Genome Viewer (IGV) (58) available through the Broad Institute servers. DNA methylation 450K array data for MCF7 and endocrine resistant sublines was previously published, see Stone *et al* (59). DNA methylation of breast tumours was sourced from The Cancer Genome Atlas (TCGA) (60) and correlated to the gene expression of *MIR196A* from the TCGA cohort.

### Breast Tumour Expression Analysis

METABRIC expression and clinical information were sourced from EGAS00000083 (61, 62). Clustering of Illumina Array and miR-Seq data was performed using the Multiple Experiment Viewer (MeV, (63)). Data was mean-centred and hierarchically clustered via Manhattan average-linkage based clustering of both rows and columns. Genes were correlated within clusters using the CORREL function of Microsoft Excel.

### Survival Analysis

Univariate and multivariate Cox proportional hazard regression analyses were performed using MedCalc for Windows, version 12.7 (MedCalc Software, Ostend, Belgium). Kaplan-Meier survival analysis and generation of survival curves was done GraphPad Prism. Optimal cutoffs for low and high expression groups were determined using receiver operator characteristic (ROC) curves.

### 3C and ChIA-PET

Chromosome conformation capture (3C) was adapted from Vakoc 2005 (29), Hagege 2007 (64) and Tan-Wong 2008 (65). Briefly, cells were grown to 60-80% confluence and fixed with 1% formaldehyde. Libraries were generated for each cell line using *Hind*III with control libraries undigested and unligated, representing native gDNA without chromosome conformation. GAPDH primers (amplified fragment contains no cut sites for these enzymes) were used to determine the digestion and ligation efficiency of each library by comparing 3C-qPCR values to primers that amplify a fragment containing a *Hind*III cut site. For each 3C-qPCR, primers were designed between 100-250 bp up or downstream of each *Hind*III cut site with the primer across the putative enhancer used as bait in each 3C-qPC.

## Acknowledgements

This study makes use of data generated by the Molecular Taxonomy of Breast Cancer International Consortium. Funding for the project was provided by Cancer Research UK and the British Columbia Cancer Agency Branch (61, 62).

---

**Supplementary Figure 1**: *MIR196A* expression correlates with *HOXC* genes in breast cancer. Hierarchically clustered normalised expression for *HOXC* genes across breast tumours. Pearson correlation coefficients to MIR196A expression are indicated on the right-hand side.

**Supplementary Figure 2**: *MIR196A* is highly expressed in breast stem cells. A) Heatmap, Manhattan hierarchically clustered demonstrating expression data for HOX genes in human breast cells. Data is reads per million (RPM) for miRNAs and reads per million per kilobase (RPKM) for mRNAs. Data is log2 normalised and mean centred by row. Right, Pearson correlation coefficients for the expression of each gene again *MIR196A*. **B)** RPM for *MIR196A* across the human breast cells. Error bars represent biological replicates when available. Data for A and B sourced from GSE16368 (51). BSC = breast stem cell, BF = breast fibroblast, BME = breast myoepithelium, BLEC = breast luminal epithelial cell and HMEC = human mammary epithelial cell.

## References

1. Wilczynska A, Bushell M. The complexity of miRNA-mediated repression. Cell death and differentiation. 2015;22(1):22–33.

2. Iorio MV, Croce CM. MicroRNA dysregulation in cancer: diagnostics, monitoring and therapeutics. A comprehensive review. EMBO Mol Med. 2012;4(3):143–59.

3. Simonson B, Das S. MicroRNA Therapeutics: the Next Magic Bullet? Mini Rev Med Chem. 2015;15(6):467–74.

4. Hui AB, Shi W, Boutros PC, Miller N, Pintilie M, Fyles T, et al. Robust global micro-RNA profiling with formalin-fixed paraffin-embedded breast cancer tissues. Laboratory investigation; a journal of technical methods and pathology. 2009;89(5):597–606.

5. Hoffman AE, Zheng T, Yi C, Leaderer D, Weidhaas J, Slack F, et al. microRNA miR-196a-2 and breast cancer: a genetic and epigenetic association study and functional analysis. Cancer research. 2009;69(14):5970–7.

6. Luthra R, Singh RR, Luthra MG, Li YX, Hannah C, Romans AM, et al. MicroRNA-196a targets annexin A1: a microRNA-mediated mechanism of annexin A1 downregulation in cancers. Oncogene. 2008;27(52):6667–78.

7. Liu XH, Lu KH, Wang KM, Sun M, Zhang EB, Yang JS, et al. MicroRNA-196a promotes non-small cell lung cancer cell proliferation and invasion through targeting HOXA5. BMC cancer. 2012;12:348.

8. Sun M, Liu XH, Li JH, Yang JS, Zhang EB, Yin DD, et al. MiR-196a is upregulated in gastric cancer and promotes cell proliferation by downregulating p27(kip1). Molecular cancer therapeutics. 2012;11(4):842–52.

9. Hou T, Ou J, Zhao X, Huang X, Huang Y, Zhang Y. MicroRNA-196a promotes cervical cancer proliferation through the regulation of FOXO1 and p27(Kip1.). British journal of cancer. 2014;110(5):1260–8.

10. Wang G, Wang Y, Shen C, Huang YW, Huang K, Huang TH, et al. RNA polymerase II binding patterns reveal genomic regions involved in microRNA gene regulation. PloS one.5(11):e13798.

11. Corcoran DL, Pandit KV, Gordon B, Bhattacharjee A, Kaminski N, Benos PV. Features of mammalian microRNA promoters emerge from polymerase II chromatin immunoprecipitation data. PloS one. 2009;4(4):e5279.

12. Attema JL, Bert AG, Lim YY, Kolesnikoff N, Lawrence DM, Pillman KA, et al. Identification of an enhancer that increases miR-200b∼200a∼429 gene expression in breast cancer cells. PloS one. 2013;8(9):e75517.

13. Punnamoottil B, Rinkwitz S, Giacomotto J, Svahn AJ, Becker TS. Motor neuron-expressed microRNAs 218 and their enhancers are nested within introns of Slit2/3 genes. Genesis. 2015;53(5):321–8.

14. Bhan A, Hussain I, Ansari KI, Kasiri S, Bashyal A, Mandal SS. Antisense Transcript Long Noncoding RNA (lncRNA) HOTAIR is Transcriptionally Induced by Estradiol. \JJournal of molecular biology. 2013.

15. Milevskiy MJ, Al-Ejeh F, Saunus JM, Northwood KS, Bailey PJ, Betts JA, et al. Long-range regulators of the lncRNA HOTAIR enhance its prognostic potential in breast cancer. Human molecular genetics. 2016;25(15):3269–83.

16. Ansari KI, Hussain I, Shrestha B, Kasiri S, Mandal SS. HOXC6 Is transcriptionally regulated via coordination of MLL histone methylase and estrogen receptor in an estrogen environment. Journal of molecular biology. 2011;411(2):334–49.

17. Ansari KI, Hussain I, Kasiri S, Mandal SS. HOXC10 is overexpressed in breast cancer and transcriptionally regulated by estrogen via involvement of histone methylases MLL3 and MLL4. J Mol Endocrinol. 2012;48(1):61–75.

18. Ansari KI, Kasiri S, Hussain I, Mandal SS. Mixed lineage leukemia histone methylases play critical roles in estrogen-mediated regulation of HOXC13. The FEBS journal. 2009;276(24):7400–11.

19. Hu Z, Liang J, Wang Z, Tian T, Zhou X, Chen J, et al. Common genetic variants in pre-microRNAs were associated with increased risk of breast cancer in Chinese women. Hum Mutat. 2009;30(1):79–84.

20. Jedlinski DJ, Gabrovska PN, Weinstein SR, Smith RA, Griffiths LR. Single nucleotide polymorphism in hsa-mir-196a-2 and breast cancer risk: a case control study. Twin Res Hum Genet. 2011;14(5):417–21.

21. Siegfried Z, Eden S, Mendelsohn M, Feng X, Tsuberi BZ, Cedar H. DNA methylation represses transcription in vivo. Nature genetics. 1999;22(2):203–6.

22. Mohammed H, D’Santos C, Serandour AA, Ali HR, Brown GD, Atkins A, et al. Endogenous purification reveals GREB1 as a key estrogen receptor regulatory factor. Cell reports. 2013;3(2):342–9.

23. Hurtado A, Holmes KA, Ross-Innes CS, Schmidt D, Carroll JS. FOXA1 is a key determinant of estrogen receptor function and endocrine response. Nat Genet. 2011;43(1):27–33.

24. Magnani L, Ballantyne EB, Zhang X, Lupien M. PBX1 genomic pioneer function drives ERalpha signaling underlying progression in breast cancer. PLoS genetics. 2011;7(11):e1002368.

25. Franco HL, Nagari A, Kraus WL. TNFalpha signaling exposes latent estrogen receptor binding sites to alter the breast cancer cell transcriptome. Molecular cell. 2015;58(1):21–34.

26. Millour J, Constantinidou D, Stavropoulou AV, Wilson MS, Myatt SS, Kwok JM, et al. FOXM1 is a transcriptional target of ERalpha and has a critical role in breast cancer endocrine sensitivity and resistance. Oncogene. 2010;29(20):2983–95.

27. Heimberg A, McGlinn E. Building a robust a-p axis. Curr Genomics. 2012;13(4):278–88.

28. Sawado T, Halow J, Bender MA, Groudine M. The beta-globin locus control region (LCR) functions primarily by enhancing the transition from transcription initiation to elongation. Genes & development. 2003;17(8):1009–18.

29. Vakoc CR, Letting DL, Gheldof N, Sawado T, Bender MA, Groudine M, et al. Proximity among distant regulatory elements at the beta-globin locus requires GATA-1 and FOG-1. Molecular cell. 2005;17(3):453–62.

30. Ko JY, Oh S, Yoo KH. Functional Enhancers As Master Regulators of Tissue-Specific Gene Regulation and Cancer Development. Mol Cells. 2017;40(3):169–77.

31. Sotelo J, Esposito D, Duhagon MA, Banfield K, Mehalko J, Liao H, et al. Long-range enhancers on 8q24 regulate c-Myc. Proceedings of the National Academy of Sciences of the United States of America. 2010;107(7):3001–5.

32. Dostie J, Richmond TA, Arnaout RA, Selzer RR, Lee WL, Honan TA, et al. Chromosome Conformation Capture Carbon Copy (5C): a massively parallel solution for mapping interactions between genomic elements. Genome Res. 2006;16(10):1299–309.

33. Davies JO, Telenius JM, McGowan SJ, Roberts NA, Taylor S, Higgs DR, et al. Multiplexed analysis of chromosome conformation at vastly improved sensitivity. Nature methods. 2016;13(1):74–80.

34. Fullwood MJ, Liu MH, Pan YF, Liu J, Xu H, Mohamed YB, et al. An oestrogen-receptor-alpha-bound human chromatin interactome. Nature. 2009;462(7269):58–64.

35. Jiang CF, Shi ZM, Li DM, Qian YC, Ren Y, Bai XM, et al. Estrogen-induced miR-196a elevation promotes tumor growth and metastasis via targeting SPRED1 in breast cancer. Molecular cancer. 2018;17(1):83.

36. Landgraf P, Rusu M, Sheridan R, Sewer A, Iovino N, Aravin A, et al. A mammalian microRNA expression atlas based on small RNA library sequencing. Cell. 2007;129(7):1401–14.

37. Kron KJ, Bailey SD, Lupien M. Enhancer alterations in cancer: a source for a cell identity crisis. Genome Med. 2014;6(9):77.

38. Herz HM, Hu D, Shilatifard A. Enhancer malfunction in cancer. Molecular cell. 2014;53(6):859–66.

39. Chen H, Li C, Peng X, Zhou Z, Weinstein JN, Cancer Genome Atlas Research N, et al. A Pan-Cancer Analysis of Enhancer Expression in Nearly 9000 Patient Samples. Cell. 2018;173(2):386–99 e12.

40. Whyte WA, Orlando DA, Hnisz D, Abraham BJ, Lin CY, Kagey MH, et al. Master transcription factors and mediator establish super-enhancers at key cell identity genes. Cell. 2013;153(2):307–19.

41. Hu Y, Zhang Z, Kashiwagi M, Yoshida T, Joshi I, Jena N, et al. Superenhancer reprogramming drives a B-cell-epithelial transition and high-risk leukemia. Genes & development. 2016;30(17):1971–90.

42. Burger H. The menopausal transition--endocrinology. J Sex Med. 2008;5(10):2266–73.

43. Hale GE, Robertson DM, Burger HG. The perimenopausal woman: endocrinology and management. J Steroid Biochem Mol Biol. 2014;142:121–31.

44. Azim HA, Jr., Partridge AH. Biology of breast cancer in young women. Breast cancer research: BCR. 2014;16(4):427.

45. Azim HA, Jr., Michiels S, Bedard PL, Singhal SK, Criscitiello C, Ignatiadis M, et al. Elucidating prognosis and biology of breast cancer arising in young women using gene expression profiling. Clinical cancer research: an official journal of the American Association for Cancer Research. 2012;18(5):1341–51.

46. McClelland RA, Barrow D, Madden TA, Dutkowski CM, Pamment J, Knowlden JM, et al. Enhanced epidermal growth factor receptor signaling in MCF7 breast cancer cells after long-term culture in the presence of the pure antiestrogen ICI 182,780 (Faslodex). Endocrinology. 2001;142(7):2776–88.

47. Knowlden JM, Hutcheson IR, Jones HE, Madden T, Gee JM, Harper ME, et al. Elevated levels of epidermal growth factor receptor/c-erbB2 heterodimers mediate an autocrine growth regulatory pathway in tamoxifen-resistant MCF-7 cells. Endocrinology. 2003;144(3):1032–44.

48. Staka CM, Nicholson RI, Gee JM. Acquired resistance to oestrogen deprivation: role for growth factor signalling kinases/oestrogen receptor cross-talk revealed in new MCF-7X model. Endocrine-related cancer. 2005;12 Suppl 1:S85–97.

49. Miller DF, Yan PS, Buechlein A, Rodriguez BA, Yilmaz AS, Goel S, et al. A new method for stranded whole transcriptome RNA-seq. Methods. 2013;63(2):126–34.

50. He DX, Gu F, Gao F, Hao JJ, Gong D, Gu XT, et al. Genome-wide profiles of methylation, microRNAs, and gene expression in chemoresistant breast cancer. Scientific reports. 2016;6:24706.

51. Gascard P, Bilenky M, Sigaroudinia M, Zhao J, Li L, Carles A, et al. Epigenetic and transcriptional determinants of the human breast. Nat Commun. 2015;6:6351.

52. Welboren WJ, van Driel MA, Janssen-Megens EM, van Heeringen SJ, Sweep FC, Span PN, et al. ChIP-Seq of ERalpha and RNA polymerase II defines genes differentially responding to ligands. The EMBO journal. 2009;28(10):1418–28.

53. Hah N, Danko CG, Core L, Waterfall JJ, Siepel A, Lis JT, et al. A rapid, extensive, and transient transcriptional response to estrogen signaling in breast cancer cells. Cell. 2011;145(4):622–34.

54. Li G, Ruan X, Auerbach RK, Sandhu KS, Zheng M, Wang P, et al. Extensive promoter-centered chromatin interactions provide a topological basis for transcription regulation. Cell. 2012;148(1-2):84–98.

55. Ross-Innes CS, Stark R, Teschendorff AE, Holmes KA, Ali HR, Dunning MJ, et al. Differential oestrogen receptor binding is associated with clinical outcome in breast cancer. Nature. 2012;481(7381):389–93.

56. Langmead B, Trapnell C, Pop M, Salzberg SL. Ultrafast and memory-efficient alignment of short DNA sequences to the human genome. Genome biology. 2009;10(3):R25.

57. Zhang Y, Liu T, Meyer CA, Eeckhoute J, Johnson DS, Bernstein BE, et al. Model-based analysis of ChIP-Seq (MACS). Genome biology. 2008;9(9):R137.

58. Robinson JT, Thorvaldsdottir H, Winckler W, Guttman M, Lander ES, Getz G, et al. Integrative genomics viewer. Nature biotechnology. 2011;29(1):24–6.

59. Stone A, Zotenko E, Locke WJ, Korbie D, Millar EK, Pidsley R, et al. DNA methylation of oestrogen-regulated enhancers defines endocrine sensitivity in breast cancer. Nat Commun. 2015;6:7758.

60. Cancer Genome Atlas N. Comprehensive molecular portraits of human breast tumours. Nature. 2012;490(7418):61–70.

61. Curtis C, Shah SP, Chin SF, Turashvili G, Rueda OM, Dunning MJ, et al. The genomic and transcriptomic architecture of 2,000 breast tumours reveals novel subgroups. Nature. 2012;486(7403):346–52.

62. Dvinge H, Git A, Graf S, Salmon-Divon M, Curtis C, Sottoriva A, et al. The shaping and functional consequences of the microRNA landscape in breast cancer. Nature. 2013;497(7449):378–82.

63. Eisen MB, Spellman PT, Brown PO, Botstein D. Cluster analysis and display of genome-wide expression patterns. Proceedings of the National Academy of Sciences of the United States of America. 1998;95(25):14863–8.

64. Hagege H, Klous P, Braem C, Splinter E, Dekker J, Cathala G, et al. Quantitative analysis of chromosome conformation capture assays (3C-qPCR). Nature protocols. 2007;2(7):1722–33.

65. Tan-Wong SM, French JD, Proudfoot NJ, Brown MA. Dynamic interactions between the promoter and terminator regions of the mammalian BRCA1 gene. Proceedings of the National Academy of Sciences of the United States of America. 2008;105(13):5160–5.

